# Influence of an Exoskeleton on Inter and Intra Task Cycle Variations in Muscle Demands During Intermittent Bending

**DOI:** 10.1101/2025.09.08.674675

**Authors:** Pranav Madhav Kuber, Behnam Kazempour, Mohammad Mehdi Alemi, Ehsan Rashedi

## Abstract

Temporal effects of exoskeletons can be effectively understood through assessments that incorporate realistic conditions, while using accurate instrumentation in controlled settings. We tracked variations in back and leg muscle activity as twelve participants performed intermittent task cycles involving symmetric and asymmetric bending with a Back-Support Industrial Exoskeleton (BSIE). Comparative assessments were conducted within and across multiple task cycles with fatigue progression. BSIE offered benefits in reducing back demands (9-22%) during bending portion. However, the device increased activity during retraction (80-11%). During sustained bending, asymmetry reduced BSIE benefits in right back region (13% vs. 22% in symmetric postures) and larger benefits of 11-15% were seen in back muscles at low vs. medium fatigue levels. Recommendations indicate having a ratio of dynamic: sustained activities < 30:1 to ensure benefits of BSIEs in back. Overall, outcomes can be helpful for understanding temporal benefits of BSIEs for trunk bending tasks.

## 1. Background

Industrial Exoskeletons (EXOs) have shown promise as an effective ergonomic intervention for mitigating overexertion injuries in industrial tasks by reducing muscle demands. Their effects on the human body have been quantified through numerous controlled laboratory assessments involving bending (Bosch et al., 2016a; Kang and Mirka, 2023a) and lifting tasks (Poliero et al., 2021; Schmalz et al., 2022). Such evaluations consist of assessing variations in subjective (e.g., perceived exertion level) and objective (e.g., activation in muscle groups) measures of physical demands. Findings from such comparative evaluations can then be used to determine benefits/limitations of using the assistive device. These wearable assistive devices are usually classified based on the body region they support (upper-body (Iranzo et al., 2020) and lower-body (Onofrejova et al., 2022). Back-Support Industrial Exoskeletons (BSIEs) are a type of upper-body EXOs that aim to reduce low-back muscle demands, and ultimately the low-back injury risk when performing prolonged exertion tasks (de Looze et al., 2016). BSIEs have demonstrated benefits in reducing low-back muscle demands by ~35-38% (Bosch et al., 2016a), ~56%, (Kazerooni et al., 2019), and 18-40% (Schnieders et al., 2023) during trunk bending tasks. Use of BSIEs during manual tasks requiring trunk flexion can be beneficial in reducing injury rates across industrial environments improving overall workplace safety.

Assessing muscle activity and fatigue has been known to be one of the most effective methods to evaluate physical effects of using EXOs on their wearers (Graham et al., 2009; Poon et al., 2019; Prats-boluda and Sanchis, 2022). Passive BSIEs are typically designed to decrease muscle effort per task cycle, potentially offer cumulative advantages over extended durations, thus potentially lowering fatigue rates. Fatigue is defined as “an individual’s inability to maintain the desired capabilities for a physical activity” (Edwards, 1981). Presence of fatigue can impact not only physical and cognitive abilities but also result in errors during task performance (Rosenthal et al., 2008) and increase the risk of injury (Butkeviçiüte et al., 2021). Electromyography (EMG) is an advantageous technique in tracking muscle activity patterns, where many measures such as the increase of peak amplitude or decrease of median frequency can indicate muscle fatigue (Enoka and Duchateau, 2017). Unlike calculating forces/moments using inverse dynamics modeling, EMG analysis can provide a more realistic understanding of muscular demands. Earlier studies demonstrated that using BSIE resulted in a minimal increase in peak amplitude of 26% compared to 88% increase without assistance in repetitive lifting (Lotz et al., 2009), and ~61% reductions in median frequency during sustained bending tasks (Kim et al., 2024). Implications of variations in such measures can then be directed towards understanding the short and long-term impacts of using these assistive devices on the human body.

The effects of BSIEs are evident in laboratory studies where study participants perform a predefined task with controlled body movements (Kermavnar et al., 2021a). On the contrary, real-world tasks include diverse activities, where benefits of EXOs are reported as less prominent than lab-based evaluations (De Bock et al., 2021; Kuber et al., 2022)). For instance, muscle activity reduction in trapezius muscle with a shoulder support EXO was ~20% lower in field vs. lab study (De Bock et al., 2021). Similarly, longitudinal effects of EXOs were not observed in a field study about implementing EXOs across automotive plants (Kim et al., 2021). One possible explanation for these mixed effects could be the counterproductive nature of body movements performed within tasks involving BSIEs. Such instances may lead to adverse/side-effects like increased muscular demands and discomfort (Kranenborg et al., 2023). Given the differences observed across lab and field assessments, a more in-depth task-centric approach can be helpful to understand the contribution of common activities (static/dynamic/sustained) within task cycles to the overall benefits provided by BSIEs during trunk flexion tasks. This understanding may aid in in formulating guidelines for their appropriate usage (Schwerha et al., 2022; Stirling et al., 2020).

Most studies on passive BSIEs have focused on lifting tasks, with limited research on their effects during dynamic, intermittent bending tasks. Additionally, prior research has mainly assessed isolated static postures, missing the opportunity to capture the full complexity of real-world movement cycles. This study is an extension to our previous publication, where overall muscle activity variations with the BSIE were examined during intermittently performed bending task cycles (Kuber and Rashedi, 2025). Specifically, the current study employs a task-centric approach, conducting an in-depth spatiotemporal analysis to evaluate the impact of a BSIE both within (differences between different types of activities: static, sustained, dynamic) and across task cycles (differences over groups of multiple task cycles that were categorized based on perceived fatigue levels). Prior to experimentation, it was hypothesized that BSIE-use may reduce muscle demands during sustained activities but increase demands during dynamic activities such as bending/retraction as well as with asymmetric postures. This study aims to provide actionable insights for practitioners by analyzing how BSIEs influence different types of activities within intermittent bending cycles. Additionally, we examine the effects of fatigue on exoskeleton performance, offering a deeper understanding of their practical applications in industrial settings.

## 2. Methodology

### 2.1. Participants

Twelve adult males were recruited from university community, per the sample sizes used in prior studies, determined as sufficient to observe significant differences in muscle activity measures (Koopman et al., 2020; Poon et al., 2019). Anthropometric measurements (age, height, weight) of participant pool have been depicted in Table 1. Participant inclusion requirements were: (a) engagement in exercise no less than twice per week, and (b) absence of musculoskeletal disorder in the back/lower body within the past 6 months. Written informed consent was collected from participants prior to data collection, as approved by university’s review board (approval code: HSRO#01113021).

**Table 1:**
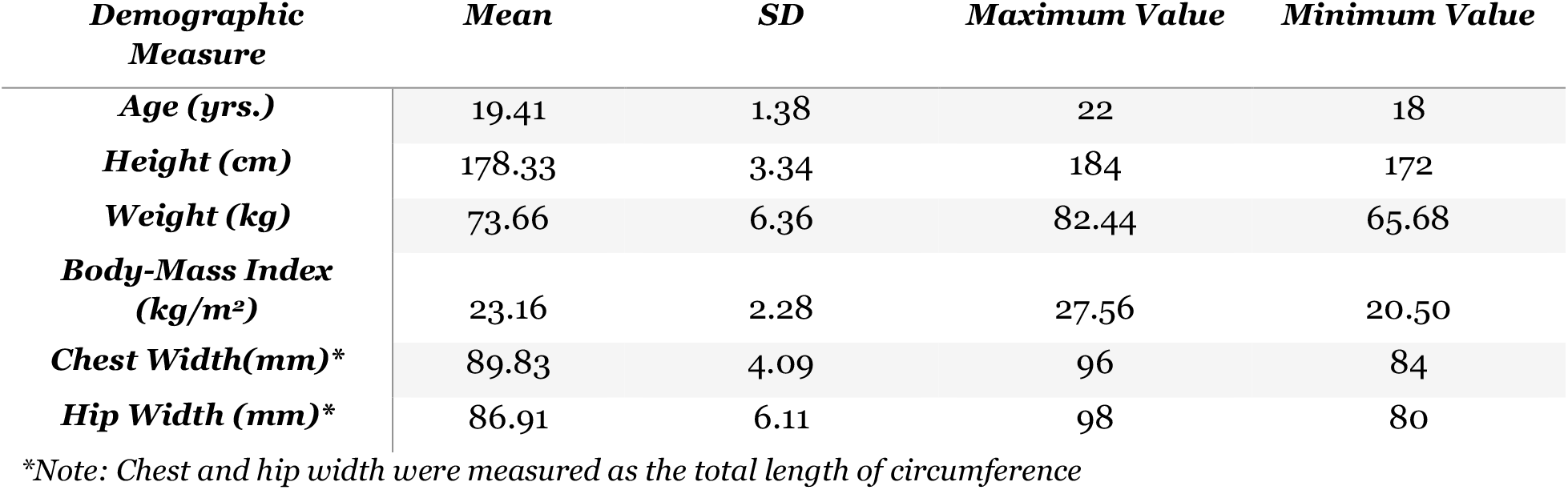
Demographic measurements of the study participants including the mean, standard deviation (SD), and ranges presented as maximum and minimum values for age, height, weight, body-mass index, chest width, and hip width.

| **Demographic Measure** | **Mean** | **SD** | **Maximum Value** | **Minimum Value** |
| --- | --- | --- | --- | --- |
| **Age (yrs.)** | 19.41 | 1.38 | 22 | 18 |
| **Height (cm)** | 178.33 | 3.34 | 184 | 172 |
| **Weight (kg)** | 73.66 | 6.36 | 82.44 | 65.68 |
| **Body-Mass Index (kg/m<sup>2</sup>)** | 23.16 | 2.28 | 27.56 | 20.50 |
| **Chest Width(mm)\*** | 89.83 | 4.09 | 96 | 84 |
| **Hip Width (mm)\*** | 86.91 | 6.11 | 98 | 80 |

\*Note: Chest and hip width were measured as the total length of circumference

### 2.2 Experimental tasks and setup

BSIEs are primarily designed to assist in trunk flexion tasks. Thus, we considered a task cycle that included static standing, dynamic bending/retraction, and sustained bending, all performed intermittently with short-duration relaxation breaks. As examining muscular demands was one of our main objectives, to ensure minimal motion-related artifacts in the collected data, a simplified wire grasping task was selected for this study wherein grasped two wire connectors placed ~10 inches apart. The wiring setup was placed on a stand with height and incline adjustments in front of the participant (adjusted to waist height of each participant such that the sternum angle was ~45 degrees in the sagittal plane). Additionally, to simulate awkward postures, an asymmetric posture were considered (Kang and Mirka, 2023b), where the setup was placed ~45 degrees in the transverse plane relative to the neutral position of the participant (Figure 1). To control trunk angles, floor markings and two height adjustable tripods with rope attachment placed in front of the participant.

**Figure 1:**
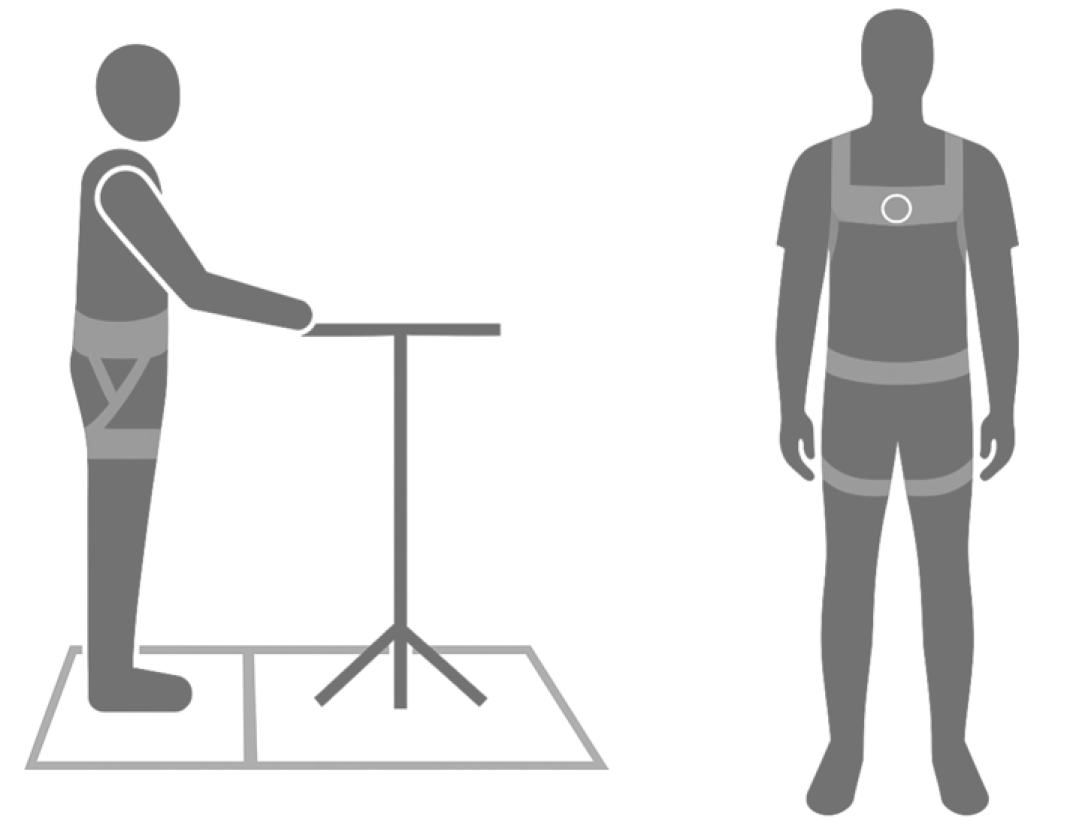
Schematic showing (left) the experimental setup, and (right) the SuitX Back-support Industrial Exoskeleton

### 2.3. Instrumentation and measurement equipment

The selected EXO was the BackX Model AC (SuitX, Emeryville, California, USA), which features a torque generation mechanism on each side of the hip region (Figure 1). We opted medium support (~25 lbs of assistance) from the three available levels (Low/Medium/High) and maintained consistency across all participants. The device was adjusted based on anthropometric measurements of participants, as instructed by the manufacturer’s manual. These adjustments were verified and fine-tuned during the familiarization session to ensure participants’ comfort.

To assess perceived exertion during task performance, participants’ perceived fatigue levels were measured using the Borg RPE CR-10 scale throughout the experiment. The scale consists of levels denoted by numbers ranging from 0-10. Levels 0, 1, 2, 3, 4, 5, 7, and 10 represent ‘no exertion’, ‘very slight’, ‘slight’, ‘moderate’, ‘somewhat severe’, ‘severe’, ‘very severe’ and ‘maximal’ levels of fatigue, respectively. Meanwhile, for objective assessment, surface EMG was used to detect electrical pulses during muscle contraction. Earlier findings indicated that time-series analysis of these pulses can provide insights into characteristics, such as the capacity of the muscle to generate force (amplitude of the signal) and muscle fatigue levels (decrease in amplitude/median frequency of signal) (González-Izal et al., 2012; Poon et al., 2019; Rashedi et al., 2014). Four EMG sensors (Delsys Trigno, Natick, Massachusetts, 1200 Hz) were attached to specific muscles using double-sided tape: two on the left/right erector spinae muscles (LES/RES), and two on left/right bicep femoris (LBF/RBF) muscles, as shown in Figure 2. The sensor placement adhered to guidelines (“SENIAM project (Surface ElectroMyoGraphy for the Non-Invasive Assessment of Muscles). Trunk movement was recorded using IMU (Inertial Measurement Unit) sensors integrated into each EMG sensor.

**Figure 2:**
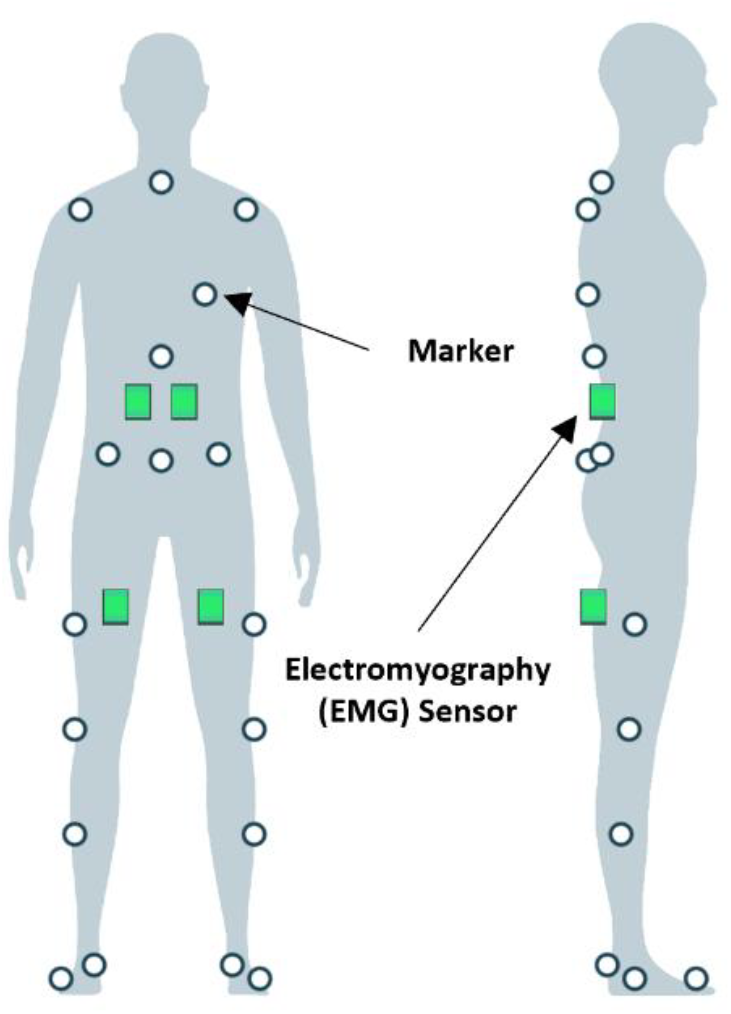
Schematic depicting sensor placement for data acquisition, including surface electromyography sensors placed on erector spinae and biceps femoris muscles.

### 2.4. Experimental design and procedure

This study consisted of three sessions, each separated by approximately 48 hours to allow for muscle recovery from fatigue. The training session (session_0) focused on familiarizing participants with the BSIE, experimental procedures as well as the Borg scale. Subjective ratings were calibrated at the beginning of the training by asking participants to perform a wall-sit task. They verbally expressed their perceived exertion ratings using the Borg RPE CR-10 scale until reaching maximal fatigue. Subsequently, EMG sensors were affixed, and Maximum Voluntary Contractions (MVCs) were performed to isolate specific muscles (LES, RES, LBF, RBF) by manually restricting body movement (Figure 3).

**Figure 3:**
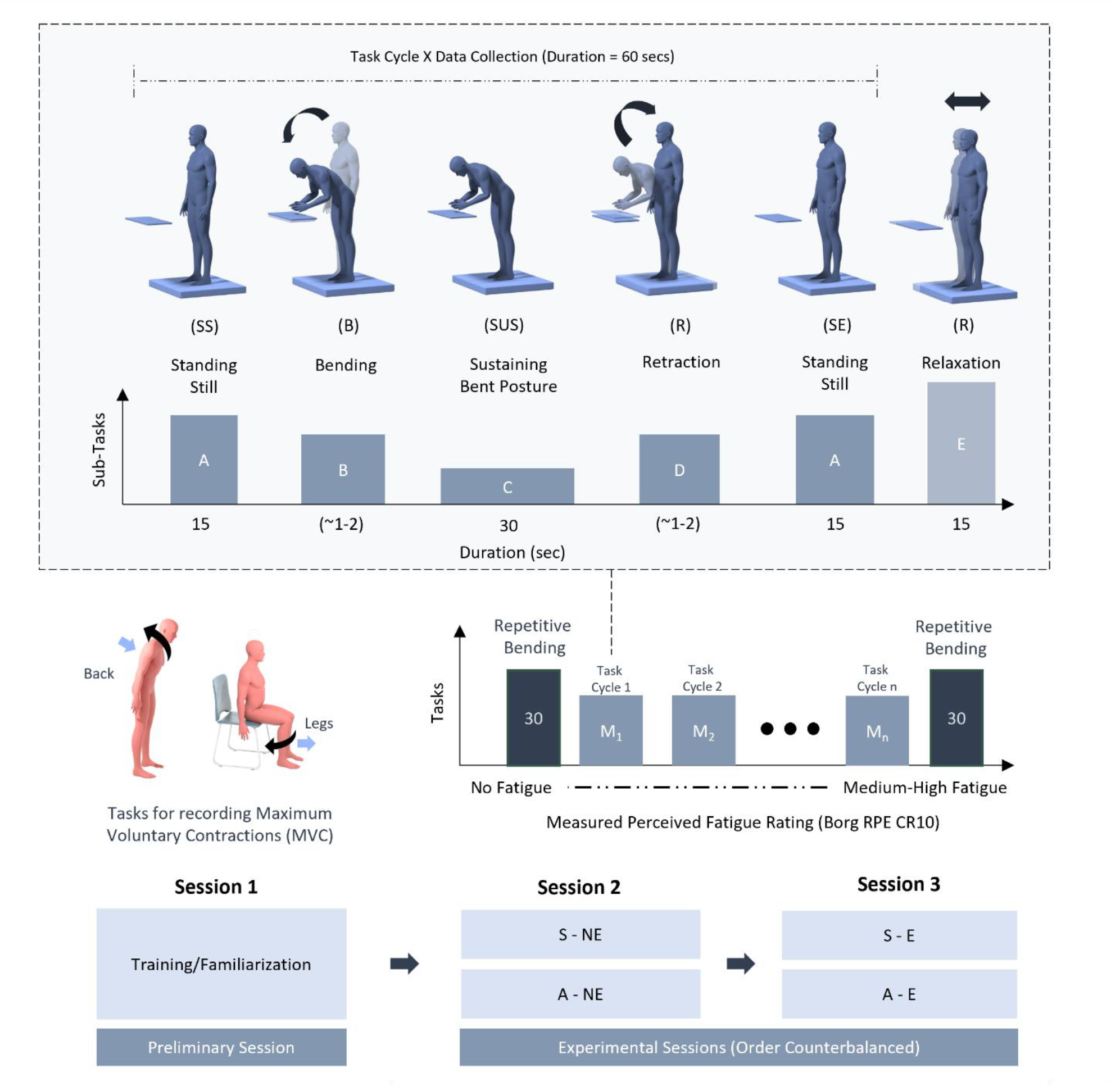
Schematic depicting the experimental protocol, tasks and activities within each task cycle.

A 2 × 2 experimental design featured independent variables as Assistance (No EXO/NE, BSIE/E) and Posture (Symmetry/S, Asymmetry/A). This led to a total of four conditions (S-E, S-NE, A-E, A-NE). The session_0 involved participants performing tasks in all four conditions, familiarizing themselves with the BSIE through walking and bending activities. After completing task cycles for the first condition, participants took a rest break until reaching medium high fatigue level (7) in the back region on the Borg scale. Each NE session lasted ~2-3 hrs. (including S and A conditions), while each E session took ~4-5 hrs. (Figure 3). Rest breaks between S/A conditions lasted ~15-25 minutes. The protocol for each condition, as illustrated in Figure 3, comprised 30 cycles of repetitive bending at the start and end, with intermittent bending modules in between. Each module included standing in a neutral posture (15 secs), sustaining a bent posture (30 secs), standing still (15 sec), and relaxing (15 sec). These durations were determined through pilot studies, considering fatigue levels and session completion times. Participants could move/rotate their torso and limbs during relaxation, providing exertion ratings on the BORG CR-10 scale. Data was recorded for each task cycle (60-sec period × 709 task cycles), totaling ~11 hrs. Following the completion of all tasks, participants rested for 10 minutes before concluding the session.

### 2.5. Data pre-processing, analysis, statistical analysis

Raw objective data were exported from the Nexus software (VICON, Hauppauge, NY) into Excel files, each representing one complete task cycle (duration: 60 secs). Each file included combined data from the EMG, force plate, and motion capture systems. Task portions were segmented based on the acceleration of the Inertial Measurement Unit (within the EMG sensors) from each 60-second trial into (a) static standing at start, (b) bending, (c) sustained: maintaining a bent posture, (d) retraction, and (e) static standing at the end. During segmentation, the middle 20 seconds of sustained bent posture, 10 seconds of standing at the start and end, and 5-95% of bending/retraction movement were selected to reduce the effects of potential movements/fluctuations.

A custom-written code was developed to initially segregate data from different systems and subsequently preprocess it. EMG data underwent filtering using a Butterworth filter (30–300 Hz, bidirectional). Then, (Bosch et al., 2016b; Chowdhury et al., 2013). For smoothing of the signals, we calculated root mean-square (RMS) envelopes using a moving window of 400 ms (Alemi et al., 2019). All EMG signals were normalized based on Peak values from MVC trials for each of the four muscles (LES, RES, LBF, and RBF). From the segmented task portions, we calculated mean values for static and sustained tasks, while peak values were derived for dynamic activities (bending/retraction). The obtained values were normalized per sensor. All objective measures for each task cycle were labeled based on the reported RPE measure, categorizing them as low fatigue (RPE <= 3) or medium fatigue (RPE >= 4). Analysis outcomes were presented as mean (SD), and statistical comparisons were conducted to assess significant differences. Post-hoc paired comparisons utilized Tukey’s Honest Significant Difference (HSD) Test where relevant. JMP Pro 14® (SAS Institute) was used for statistical analyses, with the significance level set at p-value < 0.05. Parametric model assumptions were verified before reporting results in the form of the mean (standard deviation/SD). Finally, objective measures of muscle activity in the back (LES/RES) and legs (LBF/RBF) were categorized based on perceived fatigue levels (low/medium) for the back and leg regions, respectively.

## 3. Results

Statistical comparisons can be found in Appendix with *p*-values for the main (posture (P), assistance (A)) and interaction effects of posture and assistance factors (P^*^A) for static standing at start/end, dynamic bending/retraction, and sustained bending activities. All values have been also categorized according to fatigue levels into low: (RPE in the range of 0-3 and medium: RPE in range of 4-6).

The overall variations in muscle demands across activities within task cycles have been depicted in Figure 4. Highest normalized peak amplitudes (0.6 to 0.7) were observed during dynamic retraction, followed by bending activities (0.4-0.6) and sustaining bent posture (0.2-0.4), while lowest values (<0.1) were seen during static standing at the start and end in the back region. Leg muscular demands exhibited similar trends, albeit with values consistently lower than those observed in the back region. In both back and leg regions, higher values occurred during retraction than bending in back (15%) and legs (~45%). Meanwhile, on average muscle activity was ~50% lower during sustained bending compared to retraction. The benefits of BSIE were evident during bending and sustaining bent posture in the back region and during bending, sustaining bent posture, and retraction in leg region. When categorized according to back fatigue, RES activity with assistance during retraction increased by 8% (67% MVC to 73% MVC) at medium vs. low back fatigue level.

**Figure 4:**
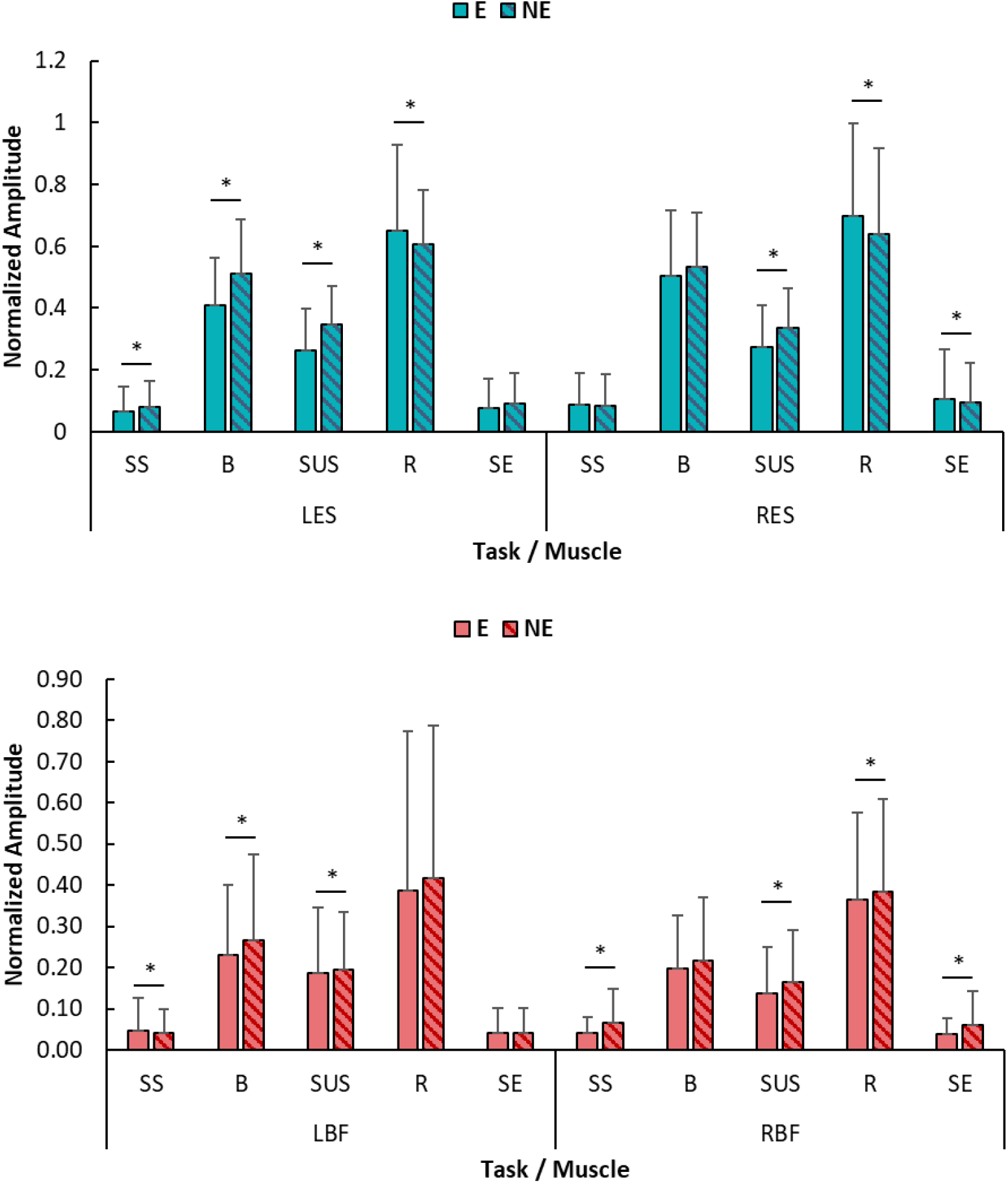
Normalized peak amplitude of (top) left/right erector spinae (LES/RES), and (bottom) biceps femoris (LBF/RBF) muscles during bending (B) and retraction (R), and mean of amplitude during standing at start (SS), sustained bending (SUS) and standing at end (SE) activities in with and without assistance (NE) conditions. (Note: *The symbol ‘*’ denotes statistical significance between E and NE conditions*)

### 3.1. Effects during Bending/Retraction

Wearing BSIE resulted in approximately 22% and 9% less activity during bending in LES (p<0.01) and RES (p<0.05) respectively, compared to not using the device (Figure 5). However, there was ~8% and ~11% more activity observed during retraction in LES (p<0.05) and RES (p=0.01) respectively. Regarding postures, lower activity was observed during both bending (~20%) and retraction (~30%) in LES while performing 45° asymmetric bending towards the left. In the leg region, BSIE led to ~12% decrease in activity in LBF (*p<*0.01) during bending. Overall, the activity was ~11% higher in LBF and ~15% lower in RBF during asymmetric vs. symmetric postures (*p<*0.01). Table 2 depicts the comparison between postures for LES and RES categorized according to back fatigue levels. RES showed lower values in asymmetric vs. symmetric postures during low fatigue levels but not during medium fatigue levels while bending as well as retraction.

**Table 2:**
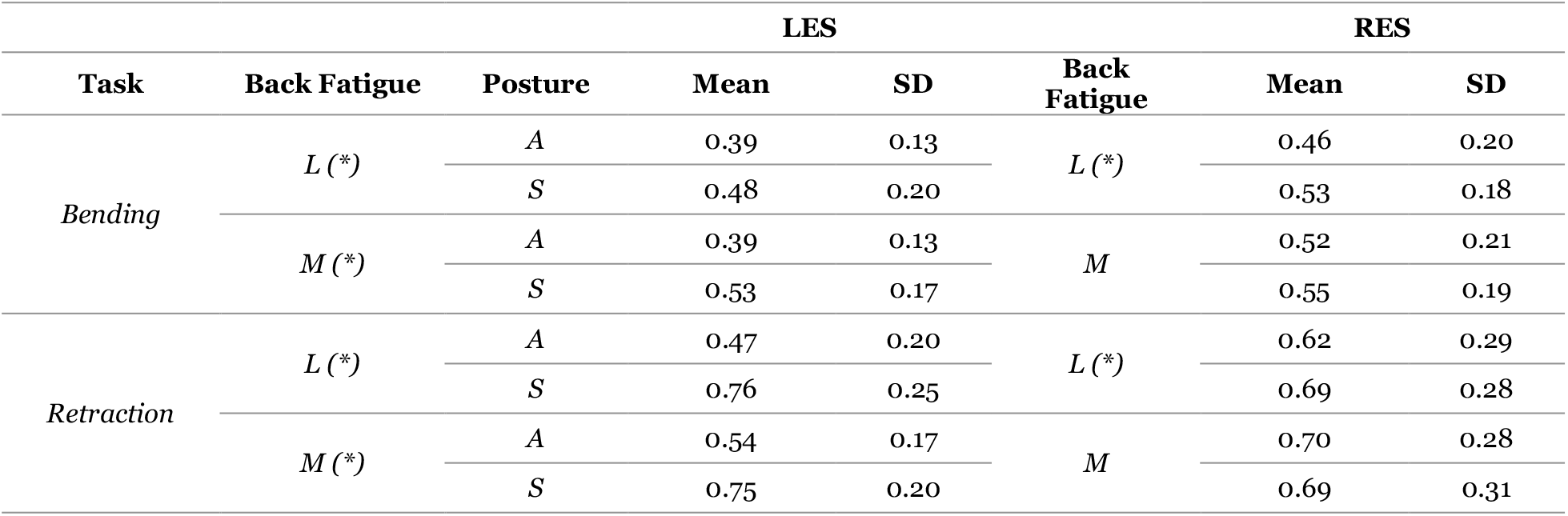
Mean (SD) values of peak amplitude in left/right biceps femoris muscles categorized according to back fatigue levels and posture. Higher values were obtained for symmetric vs. asymmetric postures (*The symbol ‘*’ denotes statistical significance between E and NE conditions*)

**Figure 5:**
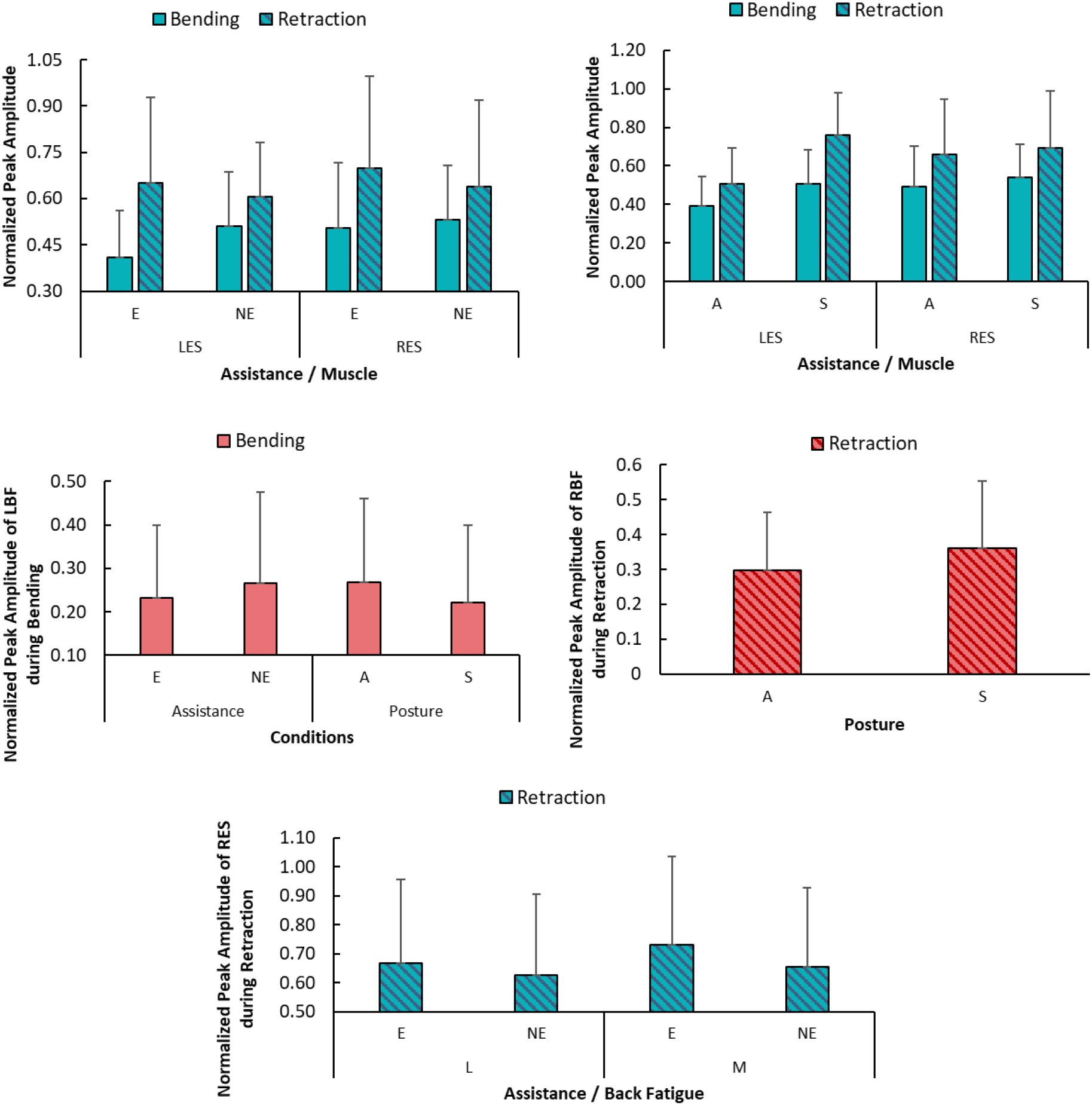
Graph showing normalized peak amplitude of left and right erector spinae muscles during bending and retraction activities across assistance (top-left) and (top-right) across postures; and (bottom-left) left biceps femoris muscle during bending across assistance and posture conditions; and (bottom-right) right biceps femoris during retraction across posture conditions. (*Note: values between levels from same factor/muscle are significant*)

### 3.2. Effects during Static Standing

The overall normalized amplitude (mean) in both back muscles (LES/RES) over static standing at the start and end portions were lower in asymmetric postures compared to symmetric postures by ~35-40% (*p*<0.05) as shown in Figure 6. Differences in RBF were seen during static standing at start and end during medium levels. RBF activity was ~59% (*p*<0.01) lower with BSIE compared to when not using assistance during medium leg fatigue (Table 3). Differences in LBF were seen at low leg fatigue levels but not at medium leg fatigue levels when categorized based on postures. In asymmetric postures, higher activity (~55%) was seen in LBF with BSIE at low fatigue levels, while lower activity (~28%) occurred in LBF with BSIE in symmetric postures (Table 4).

**Table 3:**
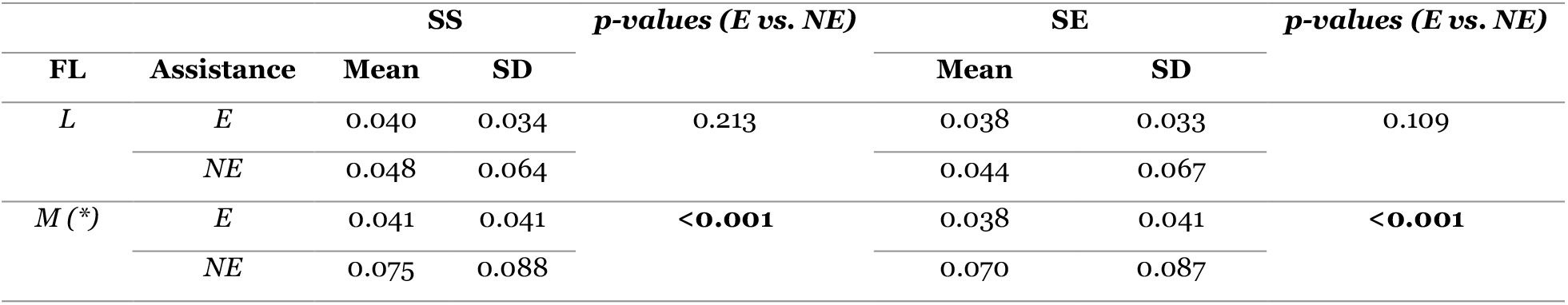
Mean (SD) values of mean amplitude in right biceps femoris (RBF) muscles categorized according to leg fatigue levels and assistance. Significant differences were seen at medium fatigue levels but not at low leg fatigue levels (the symbol ‘*’ denotes significant differences)

| FL | Assistance | SS |  | *p-values (E vs. NE)* | SE |  | *p-values (E vs. NE)* |
| --- | --- | --- | --- | --- | --- | --- | --- |
|  |  | Mean | SD |  | Mean | SD |  |
| *L* | *E* | 0.040 | 0.034 | 0.213 | 0.038 | 0.033 | 0.109 |
|  | *NE* | 0.048 | 0.064 |  | 0.044 | 0.067 |  |
| *M* (*) | *E* | 0.041 | 0.041 | <0.001 | 0.038 | 0.041 | <0.001 |
|  | *NE* | 0.075 | 0.088 |  | 0.070 | 0.087 |  |

**Table 4:** Mean (SD) values of mean amplitude in left biceps femoris (LBF) muscles categorized according to leg fatigue levels, posture, and assistance. Significant differences were seen at low fatigue levels but not at medium leg fatigue levels. (the symbol ‘*’ denotes significant difference)

| Leg Fatigue | Assistance | Posture | SS |  | SE |  |
| --- | --- | --- | --- | --- | --- | --- |
|  |  |  | Mean | SD | Mean | SD |
| *L* (*) | *E* (*) | *A* | 0.055 | 0.10 | 0.053 | 0.08 |
|  |  | *S* | 0.029 | 0.04 | 0.031 | 0.05 |
|  | *NE* (*) | *A* | 0.034 | 0.05 | 0.028 | 0.03 |
|  |  | *S* | 0.043 | 0.07 | 0.041 | 0.06 |
| *M* | *E* | *A* | 0.050 | 0.09 | 0.031 | 0.04 |
|  |  | *S* | 0.049 | 0.06 | 0.047 | 0.06 |
|  | *NE* | *A* | 0.043 | 0.05 | 0.043 | 0.06 |
|  |  | *S* | 0.046 | 0.06 | 0.044 | 0.06 |

**Figure 6:**
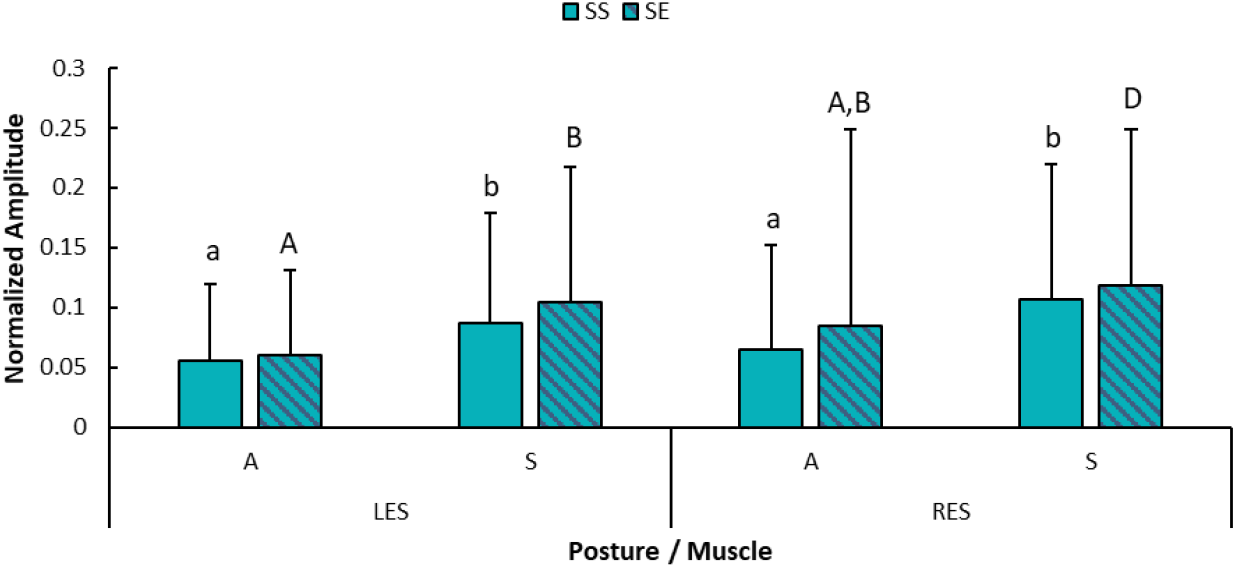
Graph showing normalized mean amplitude of left and right erector spinae (LES/RES) muscles during standing at the start (SS) and end (SE) across posture conditions. (*Note: Statistical significance is shown by different letters*)

### 3.3. Effects during Sustained Bending

The normalized mean amplitude in back muscles over sustained bending portions were lower with BSIE assistance vs. without during both asymmetric and symmetric postures. LES and RES activity were approximately 22% and 13% lower (p<0.01) with BSIE compared to without BSIE during asymmetric posture. Similarly, during symmetric posture, LES and RES activity were 25% and 22% lower (p<0.01) with BSIE compared to without BSIE (Figure 7 (top)). When categorized according to back fatigue levels, both LES and RES activity showed reduction with BSIE vs. without. For LES, BSIE reduced activity by 33% (*p*<0.01) during low back fatigue. However, this reduction decreased to 22% (*p*<0.01) during medium back fatigue. For RES, the reduction was 27% (*p*<0.01) during low back fatigue. This translated to 12% (*p*<0.01) during medium back fatigue levels (Figure 7 (bottom)). Meanwhile, in leg muscles slightly lower values with BSIE vs. without were seen during low leg fatigue in LBF (16%, *p*<0.05) and in medium fatigue levels (12%, *p*=0.01) in RBF (Table 5).

**Table 5:**
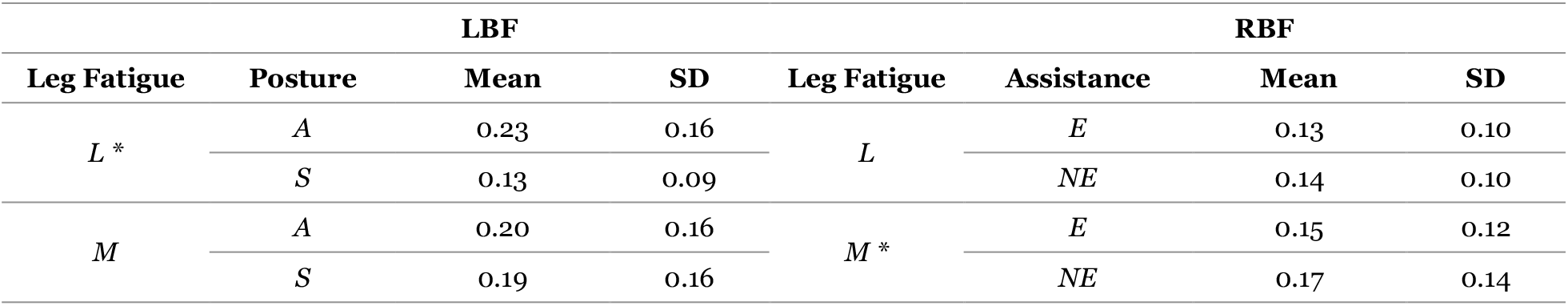
Mean (SD) values of mean amplitude in left/right biceps femoris muscles categorized according to leg fatigue levels and posture for LBF and assistance for RBF. (*significant differences are shown by ‘*’*)

| LBF |  |  |  | RBF |  |  |  |
| --- | --- | --- | --- | --- | --- | --- | --- |
| Leg Fatigue | Posture | Mean | SD | Leg Fatigue | Assistance | Mean | SD |
| $L^*$ | $A$ | 0.23 | 0.16 | $L$ | $E$ | 0.13 | 0.10 |
| | $S$ | 0.13 | 0.09 | | $NE$ | 0.14 | 0.10 |
| $M$ | $A$ | 0.20 | 0.16 | $M^*$ | $E$ | 0.15 | 0.12 |
| | $S$ | 0.19 | 0.16 | | $NE$ | 0.17 | 0.14 |

**Figure 7:**
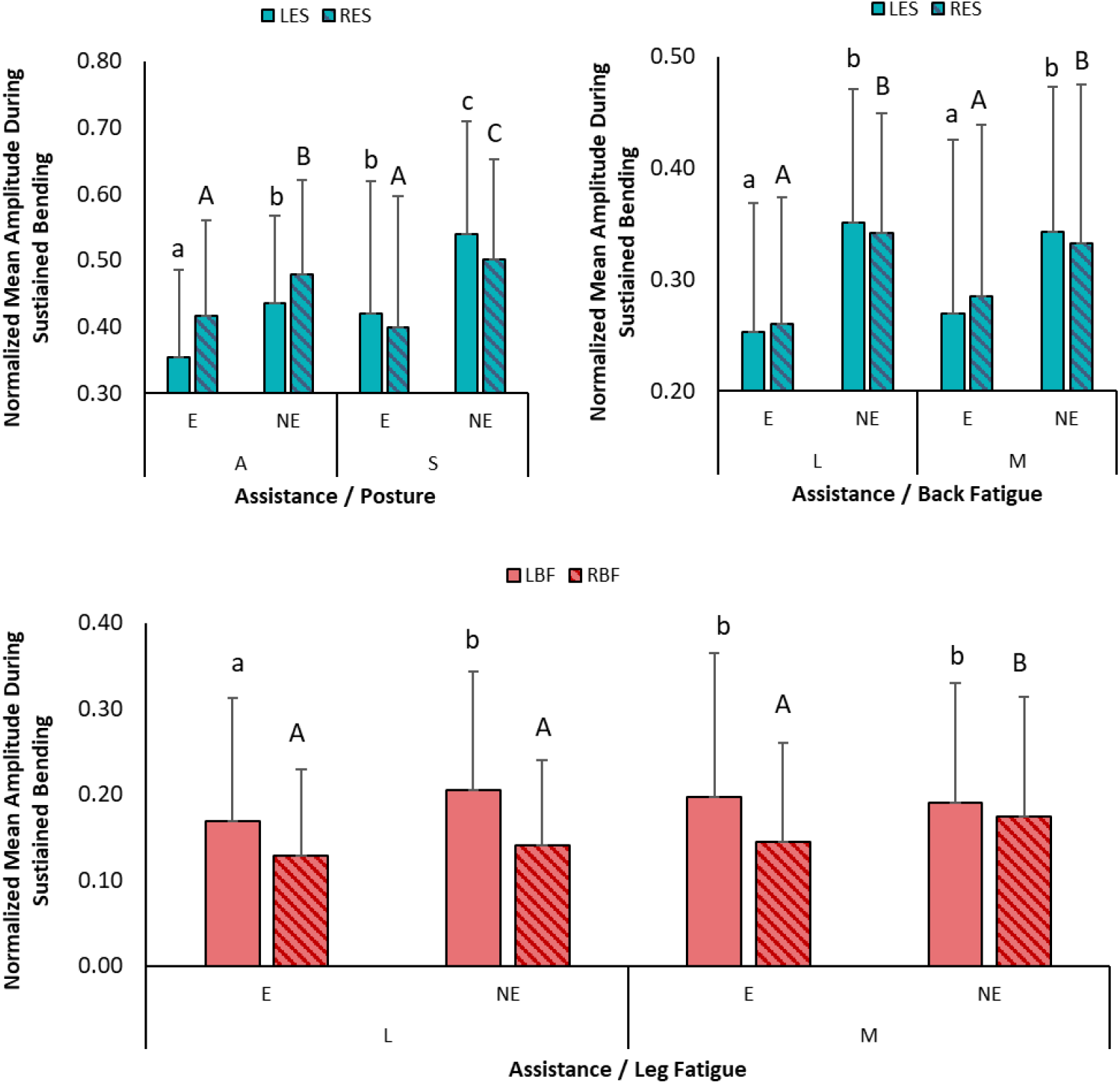
Graph showing normalized mean amplitude of left and right erector spinae (LES/RES) muscles during sustained bending across posture and assistance conditions. (*Statistical significance is shown by different letters*)

## 4. Discussion

This study evaluated the impacts of a back-support exoskeleton (BSIE) on low-back and leg muscle demands during intermittent trunk bending task, using a design that integrates elements of controlled lab experiments with the variability of real-world scenarios. By segmenting complex task cycles into distinct activity types, we were able to isolate how muscle demands vary not only across movement phases but also as fatigue progresses. This structure allowed us to bridge gaps between existing research in lab-based (Bosch et al., 2016a; Kang and Mirka, 2023a), realistic tasks in lab-spaces (Luger et al., 2023), and field studies with uncontrolled tasks (Di Natali et al., 2024).

### 4.1. Intra Task Cycle Variations

We assessed variations in muscle activation within each task cycle, in addition to evaluating overall effects of BSIE during trunk flexion. Among these activities, BSIE showed benefits during bending (LES: ~22% (*p<*0.01), RES: ~9% (*p<*0.05)) but increased demands during retraction (LES: ~8% (*p<*0.05), RES: ~11% (*p<*0.01)) (Figure 5). Similar activity reduction of ~9-12% was observed in one recent study at trunk flexion ranges of 30-60° as our study (Schwartz et al., 2021). During retraction, low-back muscles are fully contracted to pull the entire weight of the torso, as also depicted by highest activation (60-70% MVC) occurring during this portion of the task cycles. While wearing the BSIE, the wearers are required to pull the added weight of the device (~3.6 kg) which increased back activity. When performing asymmetric bending towards the left, overall, LES activity was ~20% and ~30% less during bending and retraction respectively as compared to symmetric tasks. Lesser demands in LES muscle were expected as higher contraction occurred in RES muscle bearing greater demands. Similar activity observed in RES during asymmetric and symmetric postures also supports this scenario. Similar outcomes have been reported in a recent study (Kang and Mirka, 2023b), where activity in contralateral side was higher than ipsilateral ES muscles by ~2% during 30° asymmetric bending. During bending, 11% higher activity occurred in LBF, but was ~15% lower in RBF in asymmetric vs. symmetric postures (*p<*0.01). This indicated that bending towards the left potentially resulted in higher ground reaction forces (GRF) at the left foot, increasing demands on the left leg while simultaneously decreasing demands on the right leg. We observed lower values in back muscles during static standing at the start and end portions during asymmetric vs. symmetric postures by ~35-40% (*p*<0.05) (Figure 6). These elevated values may have resulted from the increased demands observed during other phases of the task cycles, such as bending, retraction, and sustained bending, where higher demands were observed during symmetric bending.

During sustained bending, BSIE reduced LES and RES activity by ~22% and ~13% lower (*p*<0.01) during asymmetric posture, and higher reductions of 25% and 22% lower (*p*<0.01) occurred during symmetric posture (Figure 7). Hence, it could be concluded that the BSIE was particularly advantageous when sustained bending activity did not involve asymmetric postures towards the left, especially for the right muscle. This was not the case in a prior study, where asymmetry did not affect the benefits provided by the BSIE, which could be attributed to their BSIE being in the form of a soft exosuit vs. a rigid EXO in our study (Kang and Mirka, 2023b). Earlier studies demonstrated low-back reductions of 35-38% (Bosch et al., 2016a), ~56% (Kazerooni et al., 2019), and 16-34% (Schnieders et al., 2023) in ES muscles. However, it should be noted that these studies primarily assessed effects of BSIE during sustained holding postures. Fatigue increases peak amplitude of RMS of the signal. On the contrary, the rest breaks implemented in our study resulted in a delayed onset of muscle fatigue in both the back and legs, potentially offering a more realistic depiction of industrial tasks that incorporate work-rest ratios for workers. “The greater percentage reductions observed in previous studies may also stem from lower normalized values, likely influenced by higher percentage of maximum voluntary contraction (%MVC) values. For example, (Schnieders et al., 2023) reported values of 8-17% MVC values without BSIE that reduced to 6-13% with BSIE. In our study, we enrolled young adults with moderate exercise frequency and the higher %MVC values in our study (20-40% MVC) could be due to lower individual muscle capacities, or the gradual increase in peak amplitude values due to fatigue.

### 4.2. Inter Task Cycle Variations

Muscle demand variations in activities between task cycles were evaluated by compiling and categorizing measures based on perceived fatigue ratings into low (0-3) or medium (RPE: 4-6). Since fatigue progression in human musculature is highly dependent on the temporal aspects of work, low fatigue outcomes can be related to short-duration tasks, while medium fatigue outcomes can be representative of performing tasks for longer durations. An increase in peak amplitude has been previously identified as a well-known indicator of fatigue (Rampichini et al., 2020). A slower increase in EMG measures (RMS amplitude, Median frequency) at T9 and L3 joints with BSIE (~22% and 26%, 0.3% and 0.4%) compared to not using a BSIE (104% and 88%, 12% and 20%) during repetitive lifting task has been demonstrated in prior research (Lotz et al., 2009). In our study, the effect of fatigue was most clear in RES activity during retraction with assistance, wherein activity increased by 8% (67% MVC to 73% MVC) at medium vs. low back fatigue level. This was expected as low-back demands were highest during retraction phases (Figure 5), and performing activities over time increased muscle fatigue levels, especially when asymmetric postures were included.

During sustained bending portions, the BSIE provided temporal benefits in both low-back muscles, as also seen from outcomes of prior studies on BSIEs (Bosch et al., 2016a; Kermavnar et al., 2021b). Additionally, we also showed that these benefits were more pronounced at low fatigue levels compared to medium fatigue levels. For instance, for LES, BSIE reduced activity by 33% (*p<0.01*) during low back fatigue, whereas these benefits decreased to 22% (*p<0.01*) during medium back fatigue. Likewise, benefits of RES (sustained bending) were greater at low back fatigue: 27% (*p*<0.01), which reduced to 12% (*p*<0.01) during medium back fatigue levels (Figure 7 (bottom)). BSIE reduced activity in LBF (16%, *p*<0.05) during low fatigue levels. Similar benefits were seen (12%, *p*=0.01) in RBF, but during medium fatigue levels (Table 5). Mixed outcomes were found during standing at start/end portions over time. For example, BSIE reduced right leg (RBF: by ~59% (*p*<0.01)) activity during medium leg fatigue but no difference was seen during low leg fatigue (Table 3). Thus, the BSIE showed temporal benefits over longer periods of time in reducing muscle fatigue in RBF muscle. However, higher activity (10%) with BSIE vs. without was noticed in LBF during low leg fatigue levels but not at medium leg fatigue levels. This is one example of a case of redistributing loads, where the BSIE would reduce demands from one region, and increase in another; as also noted previously (Kranenborg et al., 2023). Furthermore, during low leg fatigue levels, ~55% higher activity was observed in LBF with BSIE, while ~28% lower activity occurred with BSIE in symmetric postures (Table 4). This trend could be indicating participants shifting their weight from left/right.

### 4.3. Implications on designing exoskeleton-specific task parameters

Task cycle parameters, such as number of repetitions, and duration of each activity are the key factors to determine level of exposure to physical demand. As an example, benefits from a BSIE were found to be minimal during a repetitive bending assembly task when a work-rest ratio of 1:1 was selected (stationary bending 30s, rest period 25s) (Graham et al., 2009). Similarly, duration of each activity can influence demands. In our study, we observed that during the sustained bent posture, participants experienced greater fatigue in their legs compared to their back, likely due to the consistent activation of leg muscles throughout the 60-second task cycle. On the other hand, low-back muscles were contracted mostly during the sustained bending portion (30 secs) and bending/retraction portions (~1-2 secs) of the task cycles. Using the outcomes from our evaluation, we developed an easy-to-use tool to estimate impacts of using a BSIE during bending tasks, given the parameters (duration, and number of repetitions) for each of the five activities assessed in our study, including standing still at start/end, bending, retraction, sustained bending, and relaxation (rest). A weighted means analysis was performed with means of EMG measures for each activity with/without BSIE and during asymmetric/symmetric postures along with durations as their respective weights (mean EMG amplitude during rest was taken as 1% MVC). Modified weights were then estimated by multiplying the number of repetitions to determine final weights. Lastly, weighted average was calculated using the equation:

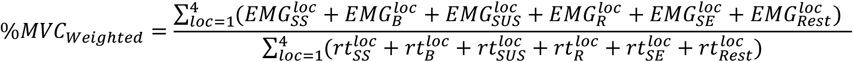

where, %*MVC*_*Weighted*_ is the estimated physical demand for task condition (AE, ANE, SE, SNE) in terms of weight average of normalized amplitude for the muscle, and (SS, B, SUS, R, SE, Rest) represent the standing at start, bending, sustained bending, retraction, standing at end, and rest periods, *loc* represents location (1:LES, 3:RES, 2:LBF, 4:RBF), and *rt* represents the modified weight obtained by multiplying duration and number of repetitions.

Overall findings for each of the experimental factors (SS/SE: 10, B/R: 3, SUS: 20, Rest: 15) have been summarized in Table 6. Using the tool, different scenarios were simulated by altering task parameters and weighted means were estimated. We averaged the outcomes for back and leg regions from the four muscles, since left/right demands may depend on minor shifts of body weight towards left/right region, possibly to relieve fatigue, as observed during experimentation. Among the simulated scenarios, outcomes showed that BSIE may be beneficial for the low-back region in tasks with higher sustained bending portions, or when rest/no back activity portions are lesser (scenario B and E in Table 6). Conversely, BSIE may not be best suited if the tasks include a higher number of bending and retraction movements. In sum, although benefits during bending (Figure 5) may compensate for increased demands during retraction while using BSIEs, our findings show that BSIE may lead higher demands if the task contains more than 30 bending and retraction movements and no sustained bending portions. More specifically, reported outcomes of this study can be used to design activities and task cycles beforehand based on task parameters. However, in some cases where industrial activities within task cycles vary highly in frequency/duration, are not well defined, and in situations where task cycles are not consistent, it may not be possible to pre-design a task cycle. In such cases, a fatigue level monitoring system, as proposed in our recent study (Kuber et al., 2024), can be more beneficial to monitor, evaluate, and intervene if there is a lack of fit between the BSIE and the task being conducted. Additionally,, as demonstrated in recent efforts towards developing adaptive EXOs (Dahmen et al., 2018; Peternel et al., 2016), findings from this study can be helpful in providing needed assistance to wearers based on type and duration of activity.

**Table 6:** Benefits of using BSIE in terms of weighted average of %MVC for each of the four conditions (A/E: Asymmetric/with Exoskeleton, S/E: Symmetric/with Exoskeleton, A/NE: Asymmetric/without Exoskeleton, S/NE: Symmetric/without Exoskeleton) during standing still at start (SS) and end (SE), sustained bending (SUS) and bending(B)/retraction (R) activities.

| Scenario | Altered Parameters | Condition | Low-Back Region |  | Thigh Region |  |
| --- | --- | --- | --- | --- | --- | --- |
|  |  |  | E | NE | E | NE |
| A | No alterations | S | 18.57 | 21.84 | 9.54 | 10.44 |
|  |  | A | 15.98 | 18.14 | 10.17 | 11.54 |
| B | Without rest, SS, and SE | S | 34.94 | 42.76 | 18.50 | 19.68 |
|  |  | A | 32.12 | 36.45 | 20.13 | 22.64 |
| C | 30 x B/R and without SUS | S | 53.43 | 52.97 | 25.72 | 27.86 |
|  |  | A | 42.60 | 44.88 | 24.57 | 26.98 |
| D | 10 x B/R and 10 secs of SUS | S | 40.43 | 41.07 | 19.62 | 21.32 |
|  |  | A | 32.51 | 34.64 | 19.05 | 21.05 |
| E | 60 secs of sustained bending, no rest/SS/SE | S | 28.68 | 38.40 | 15.86 | 16.75 |
|  |  | A | 28.05 | 32.76 | 18.17 | 20.62 |

### 4.4. Study Limitations and Future Directions

This study has several limitations that should be considered when interpreting the findings. First, participant demographics were limited to young adult males (50th percentile), which may affect the generalizability of results across different genders and body anthropometries. Future studies will explore these variations to assess potential differences in BSIE effectiveness. Second, task conditions and body movements were controlled, particularly during sustained bending phases. As fatigue progressed, participants tended to adopt a stooped posture, which may have reduced activation in key low-back muscles while increasing reliance on surrounding trunk muscles. This unintended postural shift could influence muscle demand evaluations, particularly for muscles such as the right oblique. Future studies should investigate how fatigue-induced posture adjustments affect BSIE effectiveness.

Our study categorized fatigue into two levels (low/medium), but the nonlinear recovery patterns during the relaxation phase may have influenced our findings. RPE-based assessments, while useful, are inherently subjective and may introduce self-reporting bias. Future research should incorporate more objective physiological fatigue markers to enhance measurement accuracy. Another limitation relates to fatigue progression and work-rest cycles. While we analyzed 30 cycles of repetitive bending at the start and end of each session, further investigation into how fatigue accumulates within each bending/retraction cycle could provide deeper insights. Moreover, the study utilized a work-rest ratio of 4:1 for the legs and 1:1 for the back, which may not fully reflect real-world industrial task cycles. Investigating alternative work-rest intervals could provide practical recommendations for optimizing BSIE implementation.

## Abbreviations

EXO: Exoskeleton EMG Electromyography
BSIE: Back-Support Industrial Exoskeleton RPE Ratings of Perceived Exertion
WMSD: Work-Related Musculoskeletal Disorder LES Left Erector Spinae
COP: Center of Pressure RES Right Erector Spinae
LBF: Left Biceps Femoris RBF Right Biceps Femoris
IMU: Inertial Measurement Unit

## 7. Acknowledgements

We would like to thank Dr. Maury Nussbaum for his insights, valuable guidance, and for generously sharing equipment with our lab to support our experimentation. This article is an extended version of a chapter from a doctor dissertation titled “Utilizing Exoskeletons for Intermittent Trunk Flexion Tasks: Implications on Kinematics, Stability, Muscle Activity and Fatigue”.

## 8. Conclusion

This study provides an in-depth assessment of BSIE effects on muscle demands in the low-back and leg regions during intermittent trunk flexion. By examining both inter- and intra-task cycle variations, we explored trends in muscle demands across static, dynamic, and sustained activities. Twelve participants completed 709 task cycles in symmetric and asymmetric postures, with and without BSIE assistance, progressing to a medium-high back fatigue state. A summary of the findings is shown in Table 7. BSIE effectively reduced back muscle demands during bending (LES: 22%, RES: 9%) and sustained bending (LES: ~24%, RES: ~18%), yet increased demands during the retraction phase (LES: 8%, RES: 11%). During the standing portions at the start and end of each task cycle, muscle activation shifts varied based on leg fatigue levels, likely due to asymmetric bending tasks. Similarly, in the sustained bending phase, BSIE provided leg muscle relief under certain fatigue conditions, with LBF decreasing by 16% and RBF by 12% at medium leg fatigue levels. These findings provide key insights for designing ergonomic task cycles in industrial bending environments, helping optimize BSIE use for maximum benefit. Recognizing where BSIEs offer advantages and where they may present challenges is crucial for ensuring their effective application in real-world settings. This study offers a task-centric perspective on the benefits and limitations of BSIEs, highlighting their potential to reduce musculoskeletal demands while identifying key challenges for effective real-world implementation.

**Table 7:** Summary of findings on muscle demands in low-back and legs categorized according to tasks.

| Factor | Description |
| --- | --- |
| **Overall Variation** | Order of normalized peak amplitudes (highest to lowest) in low-back:<br>60 to 70% MVC : Retraction<br>40-60% MVC : Bending<br>20-40% MVC : Sustaining bent posture,<br><0.1% MVC : Static standing at the start/end. |
| **Dynamic Bending/Retraction** | BSIE reduced activity by ~22% and ~9% in LES ($p<0.01$) and RES ($p<0.05$) during bending respectively.<br>BSIE increased activity by ~8% and ~11% during retraction in LES ($p<0.05$) and RES ($p=0.01$) respectively.<br>BSIE increased muscle activity by ~8% in RES during retraction at medium vs. low back fatigue levels.<br>BSIE led to ~12% decrease in activity in LBF ($p<0.01$) during bending but no difference was seen in RBF. |
| **Static Standing** | BSIE reduced RBF activity by ~59% ($p<0.01$) during medium leg fatigue but no difference was seen at low leg fatigue levels during both standing still portions at start and end.<br>BSIE affected LBF activity at low leg fatigue levels. In asymmetric postures, higher activity (~55%) was seen in LBF with BSIE at low fatigue levels, while lower activity (~28%) occurred in LBF with BSIE in symmetric postures (table 4). |
| **Sustained Bending** | BSIE reduced LES and RES activity by ~22% and ~13% ($p<0.01$) during asymmetric posture, and by 25% and 22% lower ($p<0.01$) during symmetric postures.<br>BSIE reduced LES activity by 33% ($p<0.01$) during low back fatigue and 22% ($p<0.01$) during medium back fatigue.<br>BSIE reduced RES activity by 27% ($p<0.01$) during low back fatigue and 12% ($p<0.01$) during medium back fatigue levels.<br>BSIE reduced activity in LBF (16%, $p<0.05$) during low leg fatigue but not during medium leg fatigue levels and (12%, $p=0.01$) in RBF during medium fatigue levels but not in low fatigue levels. |

## Appendix

Statistical comparisons between the main and interaction effects for each parameter have been shown in Table 8, while Table 9 shows the same with categorization based on perceived back and leg fatigue levels.

**Table 8:**
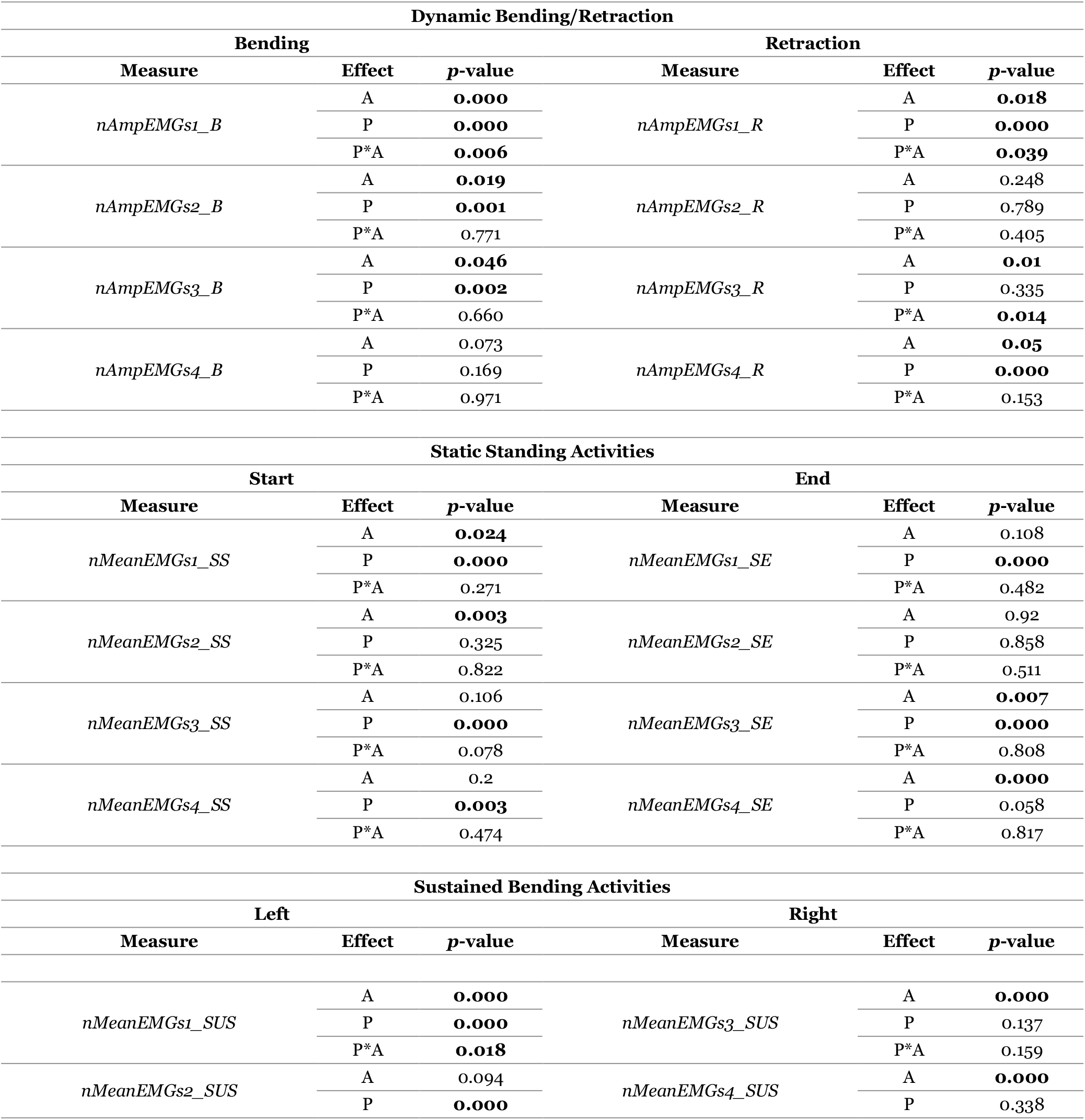
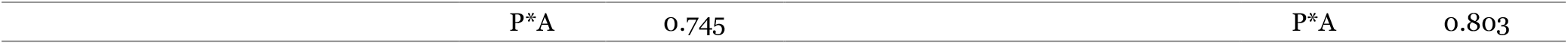
Summary statistics showing the *p*-values for peak (nAmp) and mean RMS (nMean) of left (LES: s1) and right erector spinae (RES: s3); and left (LBF:s2) and right biceps femoris (RBF: s4) muscles for the for mean and interaction effects of Assist (A), Posture Condition (P), Back Fatigue (FB), and Leg Fatigue (FL) during the dynamic activities of bending (B) and retraction (R).

| Dynamic Bending/Retraction |  |  |  |  |  |
| --- | --- | --- | --- | --- | --- |
| Bending |  |  | Retraction |  |  |
| Measure | Effect | *p*-value | Measure | Effect | *p*-value |
| *nAmpEMGs1_B* | A | **0.000** | *nAmpEMGs1_R* | A | **0.018** |
|  | P | **0.000** |  | P | **0.000** |
|  | P*A | **0.006** |  | P*A | **0.039** |
| *nAmpEMGs2_B* | A | **0.019** | *nAmpEMGs2_R* | A | 0.248 |
|  | P | **0.001** |  | P | 0.789 |
|  | P*A | 0.771 |  | P*A | 0.405 |
| *nAmpEMGs3_B* | A | **0.046** | *nAmpEMGs3_R* | A | **0.01** |
|  | P | **0.002** |  | P | 0.335 |
|  | P*A | 0.660 |  | P*A | **0.014** |
| *nAmpEMGs4_B* | A | 0.073 | *nAmpEMGs4_R* | A | **0.05** |
|  | P | 0.169 |  | P | **0.000** |
|  | P*A | 0.971 |  | P*A | 0.153 |
| Static Standing Activities |  |  |  |  |  |
| Start |  |  | End |  |  |
| Measure | Effect | *p*-value | Measure | Effect | *p*-value |
| *nMeanEMGs1_SS* | A | **0.024** | *nMeanEMGs1_SE* | A | 0.108 |
|  | P | **0.000** |  | P | **0.000** |
|  | P*A | 0.271 |  | P*A | 0.482 |
| *nMeanEMGs2_SS* | A | **0.003** | *nMeanEMGs2_SE* | A | 0.92 |
|  | P | 0.325 |  | P | 0.858 |
|  | P*A | 0.822 |  | P*A | 0.511 |
| *nMeanEMGs3_SS* | A | 0.106 | *nMeanEMGs3_SE* | A | **0.007** |
|  | P | **0.000** |  | P | **0.000** |
|  | P*A | 0.078 |  | P*A | 0.808 |
| *nMeanEMGs4_SS* | A | 0.2 | *nMeanEMGs4_SE* | A | **0.000** |
|  | P | **0.003** |  | P | 0.058 |
|  | P*A | 0.474 |  | P*A | 0.817 |
| Sustained Bending Activities |  |  |  |  |  |
| Left |  |  | Right |  |  |
| Measure | Effect | *p*-value | Measure | Effect | *p*-value |
| *nMeanEMGs1_SUS* | A | **0.000** | *nMeanEMGs3_SUS* | A | **0.000** |
|  | P | **0.000** |  | P | 0.137 |
|  | P*A | **0.018** |  | P*A | 0.159 |
| *nMeanEMGs2_SUS* | A | 0.094 | *nMeanEMGs4_SUS* | A | **0.000** |
|  | P | **0.000** |  | P | 0.338 |

|  |  |  |  |
| --- | --- | --- | --- |
| P*A | 0.745 | P*A | 0.803 |

**Table 9:** Summary statistics showing the *p*-values for the peak (nAmp) and mean RMS (nMean) values of left (LES: s1) and right erector spinae (RES: s3); and left (LBF:s2) and right biceps femoris (RBF: s4) muscles for mean and interaction effects of Assist (A), Posture Condition (P), Back Fatigue (FB), and Leg Fatigue (FL) during the sustaining bent posture (sus) portion.

| Back Fatigue (FB) Categorization of Back Demands |  |  |  | Leg Fatigue (FL) Categorization of Leg Demands |  |  |  |
| --- | --- | --- | --- | --- | --- | --- | --- |
| Measure | Effect | p-value |  | Measure | Effect | p-value |  |
|  |  | Low | Medium |  |  | Low | Medium |
| Dynamic Bending/Retraction |  |  |  |  |  |  |  |
| nAmpEMGs1_B | A | 0.000 | 0.000 | nAmpEMGs2_B | A | 0.001 | 0.473 |
|  | P | 0.000 | 0.000 |  | P | 0.000 | 0.467 |
|  | P*A | 0.001 | 0.680 |  | P*A | 0.876 | 0.719 |
| nAmpEMGs1_R | A | 0.061 | 0.197 | nAmpEMGs2_R | A | 0.003 | 0.791 |
|  | P | 0.000 | 0.000 |  | P | 0.010 | 0.118 |
|  | P*A | 0.193 | 0.104 |  | P*A | 0.774 | 0.468 |
| nAmpEMGs3_B | A | 0.071 | 0.223 | nAmpEMGs4_B | A | 0.037 | 0.568 |
|  | P | 0.001 | 0.243 |  | P | 0.984 | 0.062 |
|  | P*A | 0.772 | 0.371 |  | P*A | 0.278 | 0.769 |
| nAmpEMGs3_R | A | 0.223 | 0.027 | nAmpEMGs4_R | A | 0.083 | 0.283 |
|  | P | 0.040 | 0.05 |  | P | 0.185 | 0.000 |
|  | P*A | 0.185 | 0.522 |  | P*A | 0.387 | 0.183 |
| Static Standing |  |  |  |  |  |  |  |
| nMeanEMGs1_SE | A | 0.300 | 0.000 | nMeanEMGs2_SE | A | 0.355 | 0.319 |
|  | P | 0.000 | 0.029 |  | P | 0.966 | 0.118 |
|  | P*A | 0.059 | 0.256 |  | P*A | 0.022 | 0.176 |
| nMeanEMGs1_SS | A | 0.832 | 0.000 | nMeanEMGs2_SS | A | 0.726 | 0.527 |
|  | P | 0.000 | 0.207 |  | P | 0.624 | 0.894 |
|  | P*A | 0.067 | 0.498 |  | P*A | 0.049 | 0.754 |
| nMeanEMGs3_SE | A | 0.036 | 0.390 | nMeanEMGs4_SE | A | 0.213 | 0.000 |
|  | P | 0.011 | 0.166 |  | P | 0.104 | 0.133 |
|  | P*A | 0.881 | 0.701 |  | P*A | 0.061 | 0.178 |
| nMeanEMGs3_SS | A | 0.172 | 0.666 | nMeanEMGs4_SS | A | 0.109 | 0.000 |
|  | P | 0.000 | 0.042 |  | P | 0.097 | 0.241 |
|  | P*A | 0.014 | 0.115 |  | P*A | 0.141 | 0.176 |
| Sustained Bending |  |  |  |  |  |  |  |
| nMeanEMGs1_SUS | A | 0.000 | 0.000 | nMeanEMGs2_SUS | A | 0.013 | 0.632 |
|  | P | 0.000 | 0.000 |  | P | 0.000 | 0.228 |
|  | P*A | 0.000 | 0.309 |  | P*A | 0.434 | 0.424 |
| nMeanEMGs3_SUS | A | 0.000 | 0.004 | nMeanEMGs4_SUS | A | 0.211 | 0.010 |
|  | P | 0.725 | 0.250 |  | P | 0.051 | 0.173 |
|  | P*A | 0.001 | 0.772 |  | P*A | 0.821 | 0.842 |

